# Enzyme-constrained Metabolic Model of *Treponema pallidum* Identified Glycerol-3-phosphate Dehydrogenase as an Alternate Electron Sink

**DOI:** 10.1101/2024.11.17.624049

**Authors:** Nabia Shahreen, Niaz Bahar Chowdhury, Edward Stone, Elle Knobbe, Rajib Saha

## Abstract

*Treponema pallidum*, the causative agent of syphilis, poses a significant global health threat. Its strict intracellular lifestyle and challenges in *in vitro* cultivation have impeded detailed metabolic characterization. In this study, we present iTP251, the first genome-scale metabolic model of *T. pallidum*, reconstructed and extensively curated to capture its unique metabolic features. These refinements included the curation of key reactions such as pyrophosphate-dependent phosphorylation and pathways for nucleotide synthesis, amino acid synthesis, and cofactor metabolism. The model demonstrated high predictive accuracy, validated by a MEMOTE score of 92%. To further enhance its predictive capabilities, we developed ec-iTP251, an enzyme-constrained version of iTP251, incorporating enzyme turnover rate and molecular weight information for all reactions having gene-protein-reaction associations. Ec-iTP251 provides detailed insights into protein allocation across carbon sources, showing strong agreement with proteomics data (Pearson’s correlation of 0.88) in the central carbon pathway. Moreover, the thermodynamic analysis revealed that lactate uptake serves as an additional ATP-generating strategy to utilize unused proteomes, albeit at the cost of reducing the driving force of the central carbon pathway by 27%. Subsequent analysis identified glycerol-3-phosphate dehydrogenase as an alternative electron sink, compensating for the absence of a conventional electron transport chain while maintaining cellular redox balance. These findings highlight *T. pallidum*’s metabolic adaptations for survival and redox balance in intracellular environments, providing a foundation for future research into its unique bioenergetics.

**IMPORTANCE:** This study advances our understanding of *Treponema pallidum*, the syphilis-causing pathogen, through the reconstruction of iTP251, the first genome-scale metabolic model for this organism, and its enzyme-constrained version, ec-iTP251. The work addresses challenges of studying *T. pallidum* due to its strict intracellular nature and difficulties in *in vitro* cultivation. Validated with strong agreement to proteomics data, the model demonstrates high predictive reliability. Key insights include unique metabolic adaptations such as lactate uptake for ATP production and alternative redox-balancing mechanisms. These findings provide a robust framework for future studies aimed at unraveling the pathogen’s survival strategies and identifying potential metabolic vulnerabilities.

## INTRODUCTION

*Treponema pallidum* subspecies *pallidum* strain Nichols (hereafter *T. pallidum*), a Gram-negative spirochete responsible for syphilis, presents significant research challenges due to its strict dependence on the human host and its extreme difficulty in cultivation under laboratory conditions. Despite being identified as the causative agent of syphilis over a century ago, the continuous culture of *T. pallidum* outside a host environment remains elusive, thereby limiting investigations into how its metabolism influences its pathogenicity (1). Genome sequencing revealed a streamlined genome of approximately 1.14 million base pairs, comprising 1,041 open reading frames (2). This minimalistic architecture underscores *T. pallidum*’s heavy reliance on host-derived resources, making it one of the most metabolically reduced human pathogens known (3).

Early attempts to culture *T. pallidum in vitro* date back to 1906, following its identification as the causative agent of syphilis (4). Volpino and Fontana reported on the cultivation of the bacterium, but this initial effort faced significant challenges due to the organism’s fastidious nature and specific growth requirements (5). In 1981, Fieldsteel et al. reported the short-term multiplication of *T. pallidum* in a coculture system with rabbit epithelial cells, achieving up to 100-fold growth over 12 to 18 days (6). However, this method was not sustainable for continuous culture. A pivotal advancement occurred in 2018, when Edmondson et al. developed a refined coculture system using a modified CMRL 1066 medium supplemented with 20% fetal bovine serum, allowing *T. pallidum* to be cultivated for extended periods under controlled conditions (7). This breakthrough addressed prior challenges with contamination from rabbit epithelial cell debris in generating proteomics data, thereby facilitating an improved understanding of *T. pallidum*’s metabolic capability. Despite these advancements, implementing different synthetic biology tools (e.g., CRISPRi, gene over-expression, and gene knockouts) remains challenging in *T. pallidum*, likely due to the bacterium’s sensitivity to genetic perturbation (8). This limitation hinders functional studies of *T. pallidum*’s metabolism and virulence mechanisms.

These challenges in cultivation, coupled with its streamlined genome, underscore the need for a deeper investigation into *T. pallidum’s* metabolic strategies. Lacking a complete tricarboxylic acid (TCA) cycle, *T. pallidum* instead appears to rely primarily on substrate-level phosphorylation and acetate overflow metabolism for ATP production (9). This dependency on simplified energy generation schemes raises fundamental questions about how *T. pallidum* meets the energetic demands required to sustain its high motility—an essential virulence factor that enables tissue invasion and immune evasion (10). Recent studies suggest that *T. pallidum* supplements ATP production through additional pathways, such as a flavin-dependent acetogenic energy conservation pathway involving D-lactate dehydrogenase (D-LDH) (11). D-LDH oxidizes D-lactate to pyruvate, contributing to ATP generation. However, this mechanism introduces another question: how does *T. pallidum* regenerate the necessary reducing power to support glycolytic fluxes?

Given the complexities of studying *T. pallidum*’s metabolism and the unresolved questions about its redox strategies, we employed genome-scale metabolic models (GEMs). GEMs offer a comprehensive systems-level framework for simulating and analyzing metabolic networks, offering insights into the metabolic strategies of diverse organisms (12). Their efficacy has been demonstrated in previous studies, particularly in elucidating key phenomena in bacteria associated with sexually transmitted diseases (13, 14). Moreover, GEM can work as a unified platform to combine different ‘omics’ data to gain a holistic multi-omics understanding of an organism’s metabolism (15). This approach is especially advantageous for *T. pallidum*, where genetic perturbations remain difficult. By leveraging GEMs, we aim to provide mechanistic insights into *T. pallidum*’s metabolic adaptations, including redox-balancing strategies essential for its survival and adaptation within the host environment (16).

Accordingly, starting from an extensive literature review (1, 2, 17), we reconstructed a comprehensive GEM for *T. pallidum*, iTP251. The iTP251 achieved a MEMOTE score of 92%, demonstrating high-quality reconstruction and pathway coverage, with additional robustness assessment through FROG analysis (18, 19). To validate iTP251, we conducted *in silico* gene essentiality predictions, benchmarking them against datasets from *Escherichia coli*, a model organism with extensive essentiality data, and *Neisseria gonorrhoeae*, a human-obligate pathogen providing functional context (20, 21). This comparative validation supported the reliability of iTP251 in capturing the essential metabolic functions necessary for *T. pallidum*’s survival.

Building on this standard GEM, we developed an enzyme-constrained GEM (ecGEM), ec-iTP251, to incorporate enzyme capacity constraints not represented in traditional GEMs. While standard GEMs capture metabolism stoichiometrically, these lack representation of enzymatic limits, often leading to overestimated metabolic capacities (16, 22). The ecGEM framework allows us to integrate enzyme efficiencies and abundance, which is essential for organisms with reduced genomes like *T. pallidum*, where resource allocation is tightly regulated (13). Furthermore, the recent availability of a high-resolution proteomic dataset for *T. pallidum*, covering 94% of its proteome under *in vitro* conditions, provided an excellent benchmark for validating the ec-iTP251 (23). Comparing the model’s predicted fractional enzyme allocations with the proteomic data revealed an 88% Pearson’s correlation within central carbon pathways, reaffirming the model’s accuracy. Additionally, ec-iTP251 accurately predicted experimentally observed growth rates of *T. pallidum* for both glucose and pyruvate uptakes.

Therefore, in this study, we utilized ec-iTP251 to investigate *T. pallidum*’s unique metabolic adaptations, focusing on ATP generation schemes and redox balancing strategies. We examined how this bacterium allocates proteins and generates ATP while utilizing experimentally verified carbon sources: glucose, pyruvate, and mannose (24). Our findings revealed that protein allocation remains relatively low across these conditions, with the highest allocation observed for glucose at 29.6%, suggesting metabolic flexibility that facilitates lactate uptake. Additionally, *T. pallidum* likely employs glycerol-3-phosphate dehydrogenase as an alternative electron sink during lactate uptake, enabling redox balance and contributing to ATP production. These adaptive strategies, combined with its reliance on motility and the absence of the TCA cycle, provide selective advantages for survival and pathogenicity within the host. By highlighting these metabolic features, ec-iTP251 offers a robust platform for future studies targeting bioenergetic pathways critical for *T. pallidum*’s survival and pathogenicity.

## RESULTS AND DISCUSSIONS

### Metabolic Model Reconstruction and Refinement

To understand the metabolic landscape of *T. pallidum*, we reconstructed a genome-scale metabolic model (GEM), iTP251 (Fig. 1A). The initial draft model was generated using the KBase platform and genome data from NCBI RefSeq (KBase Genome ID: NC_021490) (25) and underwent extensive manual refinement to address key gaps in central carbon metabolism. For instance, we substituted ATP-dependent phosphofructokinase with a pyrophosphate-dependent variant and excluded the phosphotransferase system (PTS), thereby maximizing ATP utilization— a metabolic adaptation also observed in other obligate host-associated bacteria, such as *Borrelia burgdorferi* (26) (Fig. 1B).

**Fig. 1.**
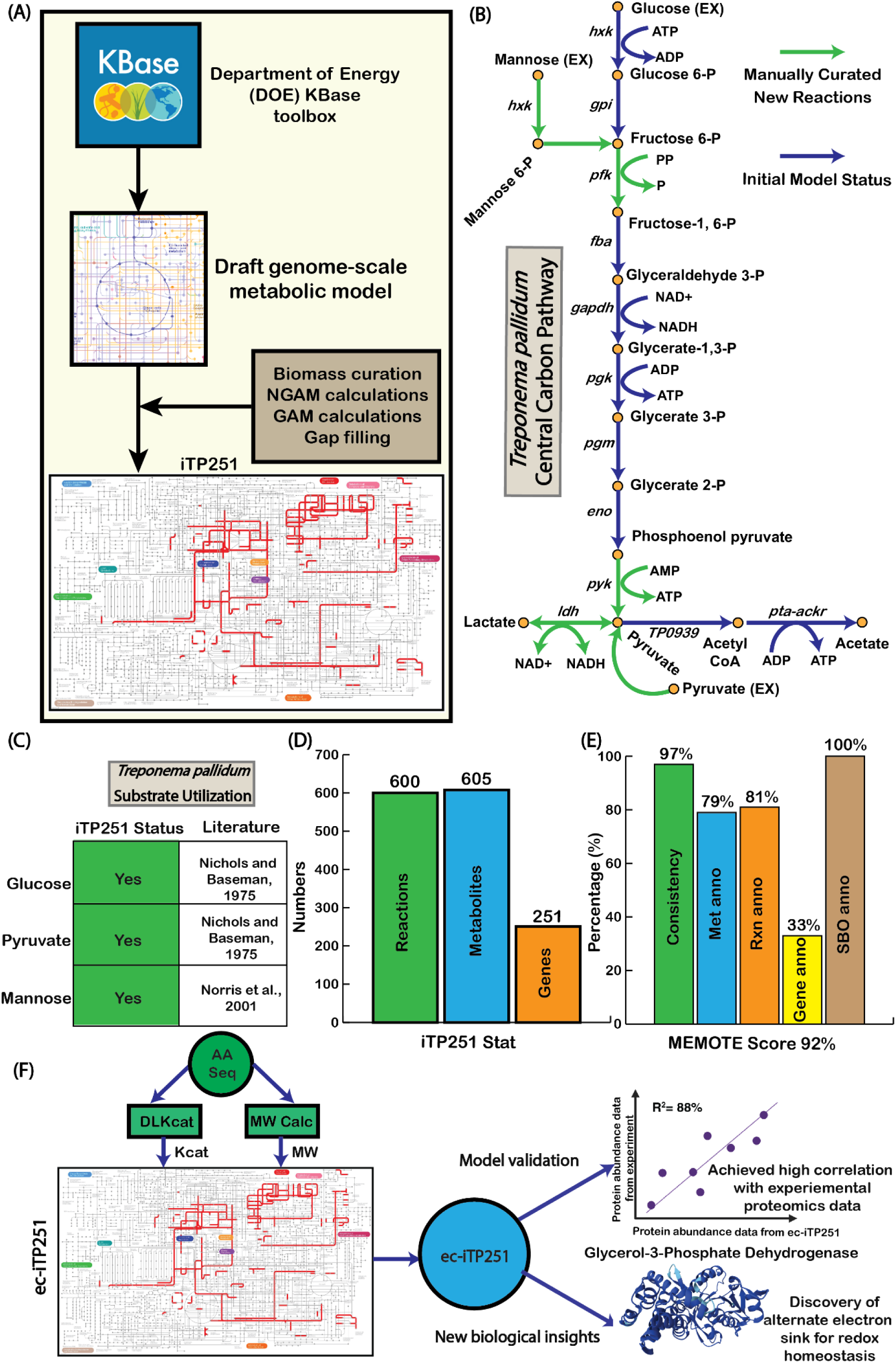
Overall process of the model reconstruction, refinement, and validation. (A) Step by step reconstruction process of iTP251 starting with the KBase platform, incorporating biomass curation, NGAM and GAM calculations, and gap filling. (B) Curated central carbon pathway showing glycolysis and key metabolic reactions, adjusted for *T. pallidum*’s unique physiology. (C) Validation of substrate utilization (glucose, pyruvate, and mannose), aligned with literature-reported metabolic capabilities (24, 27). (D) Summary of reactions, metabolites, and genes included in iTP251. (E) MEMOTE score breakdown, showcasing model consistency, annotation quality, and robustness. (F) Development and validation of the enzyme-constrained model which ultimately leads to new biological understanding (28, 29).

To address remaining gaps, we employed targeted gap-filling based on proteomic data (23), integrating essential biosynthetic and redox pathways, including those required for nucleotide, lipid, and amino acid synthesis (details in Materials and Methods). These updates allowed iTP251 to model *T. pallidum*’s growth on carbon sources that sustain its survival *in vivo*, including glucose, pyruvate, and mannose (Fig. 1C). Additionally, by incorporating D-lactate dehydrogenase (TP0037), we represent *T. pallidum*’s acetogenic ATP generation pathway, an energy-conserving adaptation in oxygen-limited environments, further distinguishing its host-adapted metabolism (11).

Regarding the biomass equation of *T. pallidum*, we took that from the phylogenetically adjacent *Borrelia burgdorferi* metabolic model, iBB151 (Table S1) (30). Using the template equation from iBB151, we reconstructed a *T. pallidum*’s specific biomass equation, integrating dry weight estimates from the literature: 70% protein, 20% lipid, and 5% carbohydrate (31). After formulating the biomass equation, we verified that the molecular weight of the biomass was 1 g/mmol (32). Additionally, growth-associated maintenance (GAM) and non-growth-associated maintenance (NGAM) values specific to *T. pallidum* were calculated as 48.69 mmol/gDW/hr and 1.50 mmol/gDW/hr, respectively, using Pirt’s equations, for capturing the energy demand of the bacterium accurately (detailed in the Materials and Methods section). Overall, iTP251 includes 600 reactions, 605 metabolites, and 251 genes (Fig. 1D).

The model’s overall quality was evaluated using MEMOTE (18), a community standard for assessing the rigor of metabolic models, resulting in an overall score of 92%, which signifies a high-quality reconstruction. Key categories, including stoichiometric and mass balance, received a score of 100%, ensuring atomic consistency across reactions, while charge balance reached 96.3%, confirming ion equilibrium within the network. Complete metabolite connectivity and 84.6% flux consistency under default medium conditions further highlight the model’s high quality. Although gene annotation coverage was 33%, reflecting gaps in *T. pallidum*’s genomic annotations from different databases, complete Systems Biology Ontology (SBO) annotation (100%) enhances transparency and compatibility with systems biology resources (Fig. 1E, Text S2).

In addition, we validated iTP251’s robustness and reproducibility using the FROG framework (33). This analysis systematically assessed flux variability, reaction deletions, objective function stability, and gene deletions, ensuring that the model’s performance remains consistent across different computational environments. The FROG report, including flux bounds, objective function values, and gene deletion data for independent reproducibility verification, is provided in Text S3.

### Validation of iTP251 Through Reaction Essentiality and Gene Essentiality Analysis

To evaluate the predictive accuracy of iTP251, we conducted comprehensive *in silico* analyses focusing on reaction and gene essentiality. Single-reaction deletions within iTP251 identified 166 essential reactions, with the majority associated with pyrimidine and purine metabolism pathways, highlighting the organism’s reliance on nucleotide synthesis (Fig. 2A). This observation aligns with prior studies that also emphasize the indispensable role of purine and pyrimidine metabolism (34). The remaining essential reactions were distributed across glycolysis and the pentose phosphate pathway, underscoring their pivotal roles in energy production and redox balance.

**Fig. 2.**
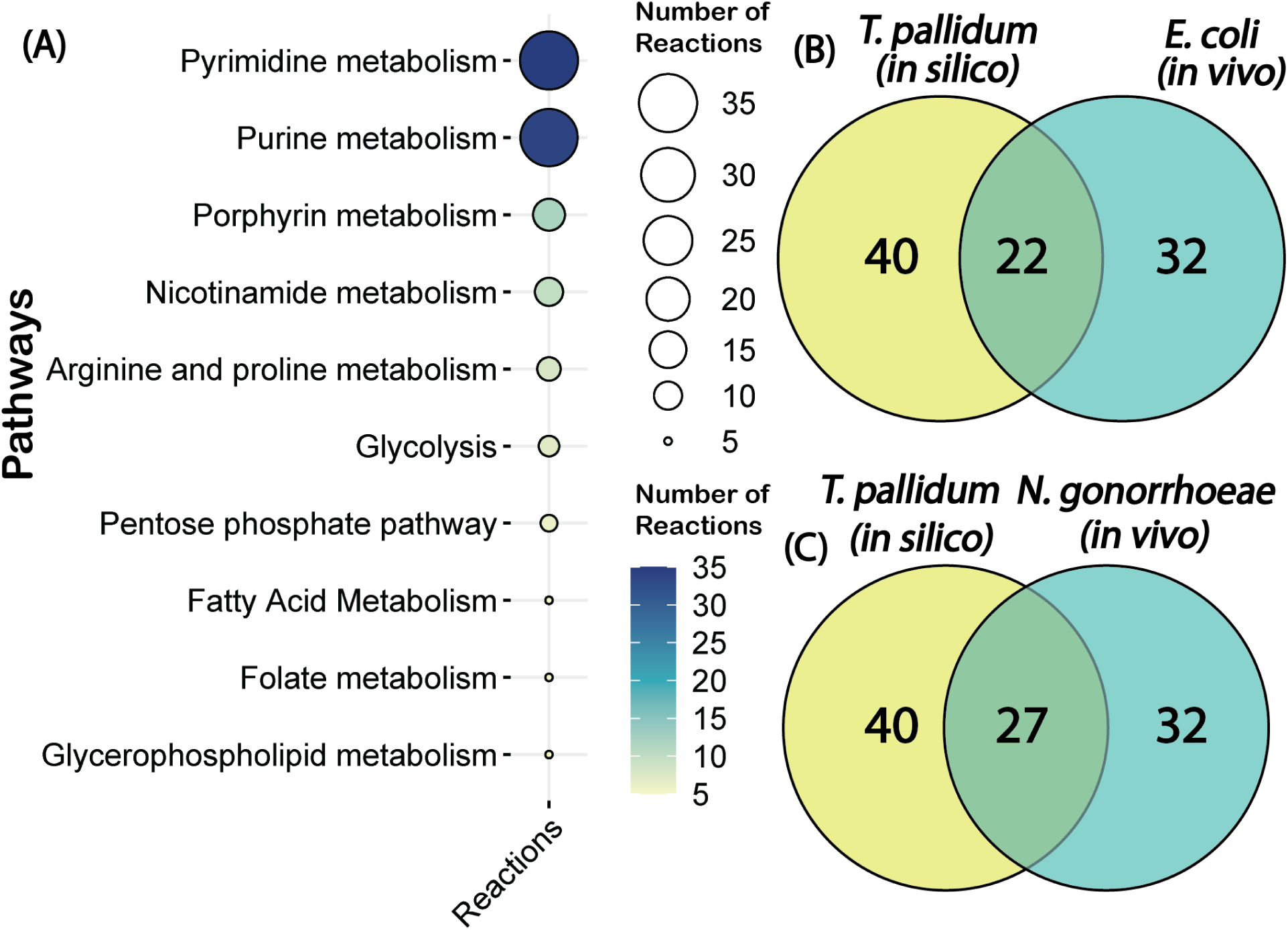
Essential metabolic pathways and comparative gene essentiality analysis. (A) Distribution of essential pathways identified in *T. pallidum* through reaction essentiality analysis, highlighting critical roles of pyrimidine metabolism, glycolysis, and the pentose phosphate pathwayl. (B) Comparative gene essentiality analysis between *T. pallidum* and *Escherichia coli*, and (C) between *T. pallidum* and *Neisseria gonorrhoeae*, underscoring conserved essential genes in Gram-negative bacteria and supporting the validity of iTP251 in representing core metabolic functions.

Of the 251 genes included in the reconstruction, 40 (15.94%) were predicted to be essential for *T. pallidum’s* survival (Table S4), a proportion comparable to other organisms (35–39). In the absence of *T. pallidum*-specific gene essentiality data, we benchmarked iTP251’s predictions against experimentally validated essential genes from *Escherichia coli* and *Neisseria gonorrhoeae*. *E. coli* was chosen for its extensive gene essentiality data as a model organism, while *N. gonorrhoeae* was selected for its relevance as an obligate human pathogen responsible for sexually transmitted infections, similar to *T. pallidum*. Among the 40 essential genes predicted by the model, 32 had orthologs in *E. coli*, with 22 experimentally confirmed as essential, yielding a concordance rate of 68.8%. Likewise, 32 orthologs were also identified in *N. gonorrhoeae*, of which 27 were experimentally validated as essential, corresponding to an 84.4% concordance rate (Fig. 2B-C). The higher concordance with *N. gonorrhoeae* likely reflects its shared traits with *T. pallidum* as a human-adapted pathogen causing sexually transmitted infections, further supporting the model’s relevance. These findings suggest that iTP251 effectively captures fundamental metabolic functions, despite potential limitations in cross-species comparisons.

### Development and Validation of the Enzyme-Constrained GEM for *T. pallidum*

Building upon iTP251, we developed an ecGEM, named ec-iTP251, to account for protein resource limitations and simulate the physiological constraints of *T. pallidum*’s metabolism under proteome-restricted conditions.

In developing ec-iTP251, we integrated enzyme turnover numbers (*K_cat_*) and molecular weights (MW) for reactions with gene-protein-reaction (GPR) associations. We utilized DLKcat (40), a computational tool leveraging sequence homology and structural properties, to predict *K_cat_* values for the majority of GPR-associated reactions. For the remaining reactions, we employed Monte Carlo simulations, generating 100 *K_cat_* samples based on the distribution of known values from the initial 391 reactions (Fig. S1). These sampled *K_cat_* values, combined with the DLKcat-derived values, were incorporated into the GEM to create 100 ensemble ecGEMs (see Materials and Method section). To identify the most representative model, we calculated the minimum cellular protein content (ϕ*_MC_*) for each model (Fig. S2) for a given growth rate and selected the one with the lowest protein demand. This process aligns with *T. pallidum*’s adaptation to protein-limited environments, consistent with its evolution as an obligate intracellular bacterium (41).

To validate ec-iTP251, we compared the model’s predicted enzyme allocations against high-resolution proteomic data encompassing 94% of *T. pallidum*’s proteome (23) (Fig. 3A). The Pearson correlation between predicted and observed fractional proteome allocations in central carbon pathways showed a strong concordance (*R*^2^ = 0.876), underscoring the model’s predictive accuracy (Fig. 3B).

**Fig. 3.**
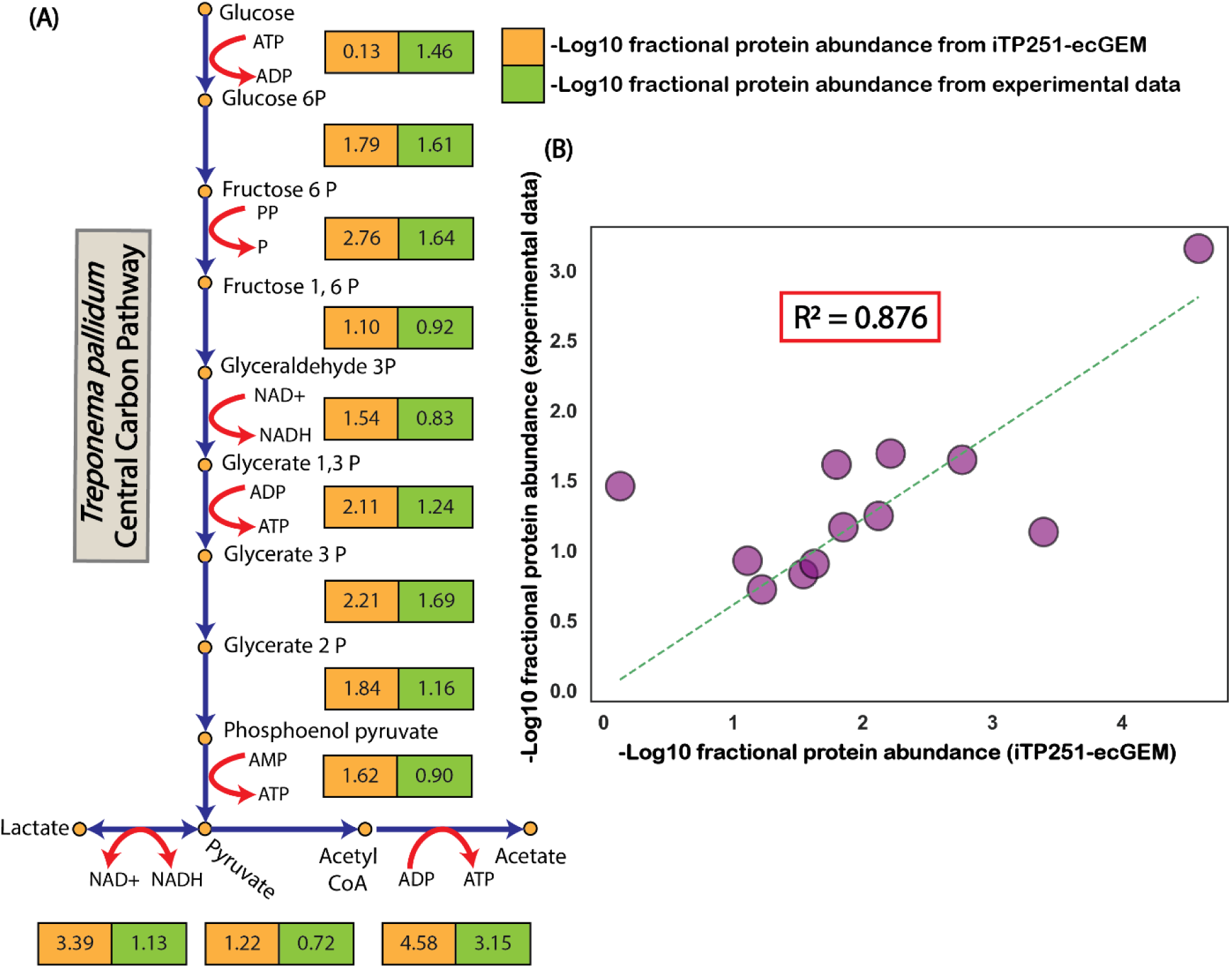
The enzyme-constrained genome-scale metabolic model (ec-iTP251) accurately predicts fractional enzyme requirements in *T. pallidum.* (A) Fractional enzyme requirements for each reaction in the central carbon pathway illustrate the distribution of protein resources across metabolic processes. (B) Correlation analysis between model-predicted fractional enzyme allocations and experimentally observed proteomic data for central carbon pathway reactions demonstrated a strong concordance (*R*^2^ = 0.876), thereby validating the predictive accuracy of ec-iTP251.

### Predicting Proteome Allocation for Different Substrates

As ec-iTP251 showed excellent accuracy in predicting the fractional enzyme abundances of the central carbon pathway, we next used ec-iTP251 to investigate *T. pallidum*’s proteomic adaptations to capture protein distribution across three experimentally validated carbon sources: glucose, pyruvate, and mannose.

When glucose was supplied as the sole carbon source, ec-iTP251 indicated a substantial allocation of proteome resources to glycolytic enzymes, reflecting *T. pallidum*’s dependence on glycolysis as a primary ATP-generating pathway (Fig. 4A). This allocation, maintained at 29.7% of total cellular dry weight, underscores the efficiency of substrate-level phosphorylation in sustaining cellular energy without overextending *T. pallidum*’s protein synthesis capacity. In contrast, when pyruvate was available, proteome resources shifted notably toward pyruvate metabolism and acetate production pathways (Fig. 4B). Due to the absence of hexokinase (*hxk)* activity, which has a very high unit flux protein cost, the protein demand for pyruvate utilization was approximately one-tenth of that for glucose, aligning with *T. pallidum*’s ability to economize proteomic resources by minimizing glycolytic flux. Pyruvate is converted directly to acetyl-CoA via pyruvate oxidoreductase, then to acetate via phosphotransacetylase and acetate kinase, which generates ATP. This metabolic shift supports “acetate overflow,” a phenomenon common in fermentative organisms that maximizes ATP output with minimal proteome investment and sustains redox balance without an electron transport chain (42). For mannose metabolism, ec-iTP251 projected an increased allocation to transporters and specialized enzymes unique to mannose uptake and processing (Fig. 4C). Despite this added investment, mannose metabolism showed proteome constraints similar to glucose, underscoring *T. pallidum*’s flexibility in utilizing alternative carbon sources available in the host environment. For glucose and pyruvate, ec-iTP251 captured the experimentally observed growth rate (Fig. 4D). For mannose, the growth rate prediction was also realistic compared to the glucose uptake (Fig. 4D).

**Fig. 4.**
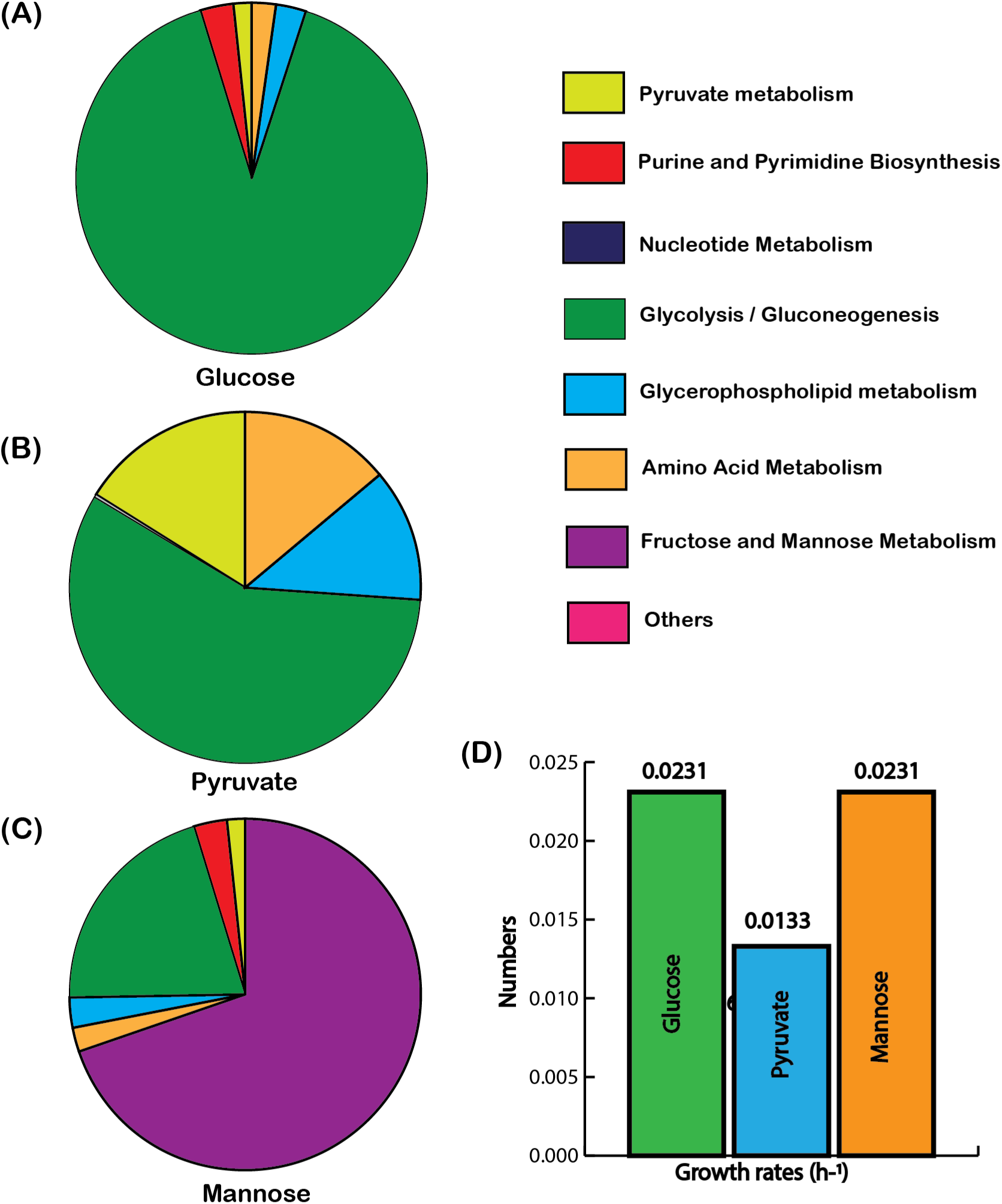
Protein allocation across major metabolic pathways for distinct carbon sources. Proteome allocation is shown for (a) glucose, (b) pyruvate, and (c) mannose as the sole carbon sources. (d) Growth rates compared across these three substrates, highlighting metabolic efficiency.

Hence, this analysis highlights *T. pallidum’s* flexibility in proteome distribution, which likely enables metabolic stability in protein-limited host environments. Unlike *E. coli*, which utilizes its complete TCA cycle and oxidative phosphorylation to distribute proteome resources more evenly across metabolic pathways (43), *T. pallidum* prioritizes glycolysis and acetate production. This streamlined proteome allocation underscores its adaptation to intracellular environments, leveraging reduced metabolic complexity to optimize ATP yield and enhance survival within the host.

### Energy Generation and Alternate Electron Sinks

After validating ec-iTP251 against proteomics data, we used the model to explore *T. pallidum*’s metabolic adaptations across the three experimentally validated carbon sources. Notably, simulation results consistently predicted lactate co-uptake alongside these substrates, revealing an alternative strategy for *T. pallidum* to generate additional ATP.

To explore the thermodynamic feasibility of this co-uptake mechanism, we applied Max-Min Driving Force (MDF) analysis to compare overall pathway driving force for lactate uptake and secretion (39) (Fig. 5A-B). The MDF analysis revealed that lactate uptake is thermodynamically 27% less favorable than lactate secretion, suggesting a need for higher protein investments to sustain lactate uptake under physiological conditions (Fig. 5C). Despite this inefficiency, the simulations suggested that lactate co-uptake supports ATP generation through TP0037, which catalyzes the oxidation of lactate to pyruvate while generating NADH. This strategy enables *T. pallidum* to utilize available proteome resources flexibly, producing additional ATP to sustain growth with minimal metabolic overhead. Furthermore, the surplus ATP generated through this pathway could be redirected to energy-intensive processes crucial for *T. pallidum*’s survival and pathogenicity, including motility and host invasion mechanisms (45).

**Fig. 5:**
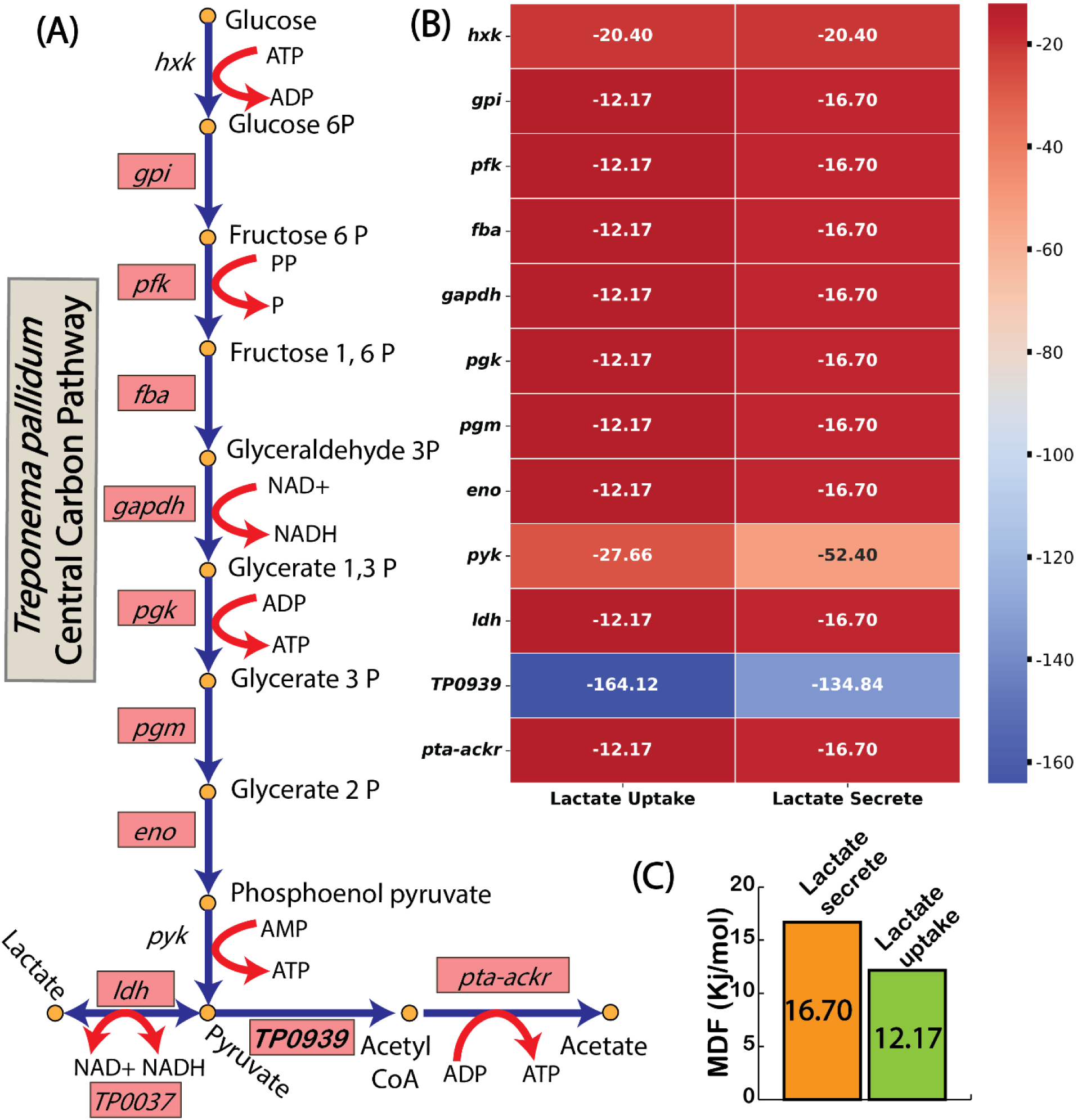
Thermodynamic constraints of lactate metabolism assessed through MDF Analysis. (A) Central carbon pathway of *T. pallidum* highlighting key reactions involved in lactate metabolism. (B) Reaction specific Gibbs free energy (Δ*G*) values during lactate uptake and secretion, showing thermodynamic feasibility. (C) Cumulative driving force analysis shows a thermodynamic preference for lactate secretion over uptake.

Given that *T. pallidum* lacks a complete TCA cycle and a functional electron transport chain; it must rely on alternative mechanisms to maintain redox balance while taking up lactate. Using the ec-iTP251 model, we investigated potential electron sinks that could sustain redox equilibrium during lactate co-uptake. Our analysis identified 16 NAD⁺ regenerating reactions within iTP251 (Table S3). Among these, three reactions demonstrated consistent non-zero flux across all tested substrates, each associated with a GPR: GPDH (glycerol-3-phosphate dehydrogenase), NADS (NAD⁺ synthase), and IspH (4-hydroxy-3-methylbut-2-enyl diphosphate reductase) (Fig. 6A). An additional NAD⁺-generating reaction was identified through gap filling but lacked a GPR association and was excluded from further analysis. Of these reactions, GPDH exhibited the highest flux, underscoring its central role as an alternative electron sink in *T. pallidum* (Fig. 6B). GPDH catalyzes the conversion of glycerol-3-phosphate to dihydroxyacetone phosphate while oxidizing NADH to NAD⁺, creating an essential pathway for NAD⁺ regeneration (Fig. 6C). This adaptation supports *T. pallidum*’s lactate uptake, facilitating additional ATP generation and redox balance without overextending its limited protein resources.

**Fig. 6.**
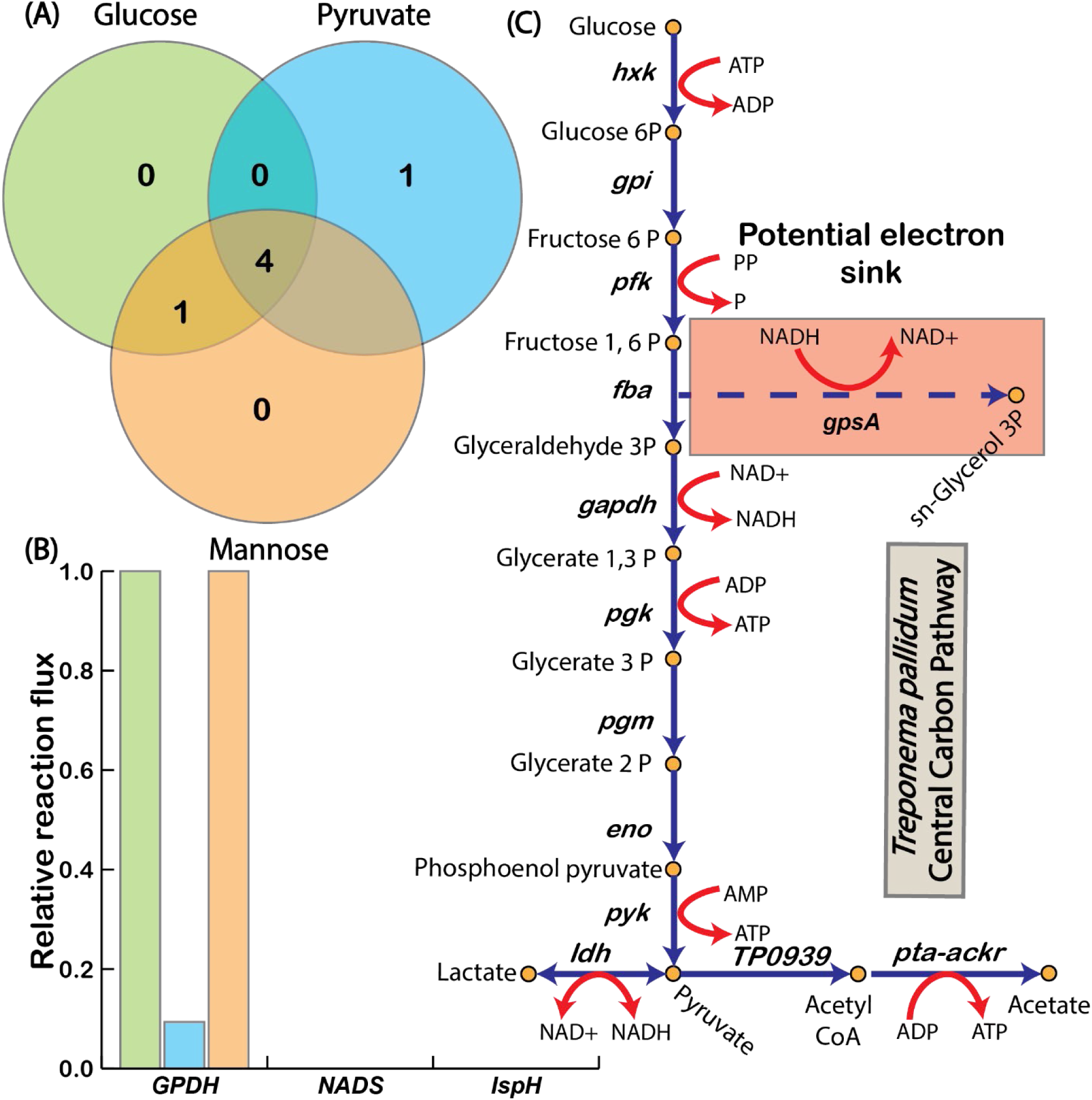
Alternate electron sink mechanisms in *T. pallidum* across different substrates. (A) Venn diagram showing shared and unique electron sinks across glucose, pyruvate, and mannose utilization. (B) GPDH carries the highest flux among alternative electron sinks, emphasizing its critical role in redox balance. (C) Schematic representation of the central carbon pathway, focusing on the *gpsA*-encoded reaction GPDH.

Interestingly, the use of GPDH as an alternative electron sink reflects strategies observed in other organisms with metabolic adaptations for redox balance. The metabolically versatile soil bacterium *Rhodopseudomonas palustris* employs GPDH to manage redox balance across diverse environmental conditions (46). Similarly, *Saccharomyces cerevisiae*, a well-studied yeast, utilizes GPDH to divert electrons toward glycerol production under anaerobic stress, bypassing the need for oxidative phosphorylation (47). These examples suggest a conserved mechanism across diverse organisms for sustaining NAD⁺ regeneration through GPDH.

In *T. pallidum*, the reliance on GPDH within a thermodynamically favorable range facilitates efficient redox balancing with minimal protein investment—an essential adaptation for an organism constrained by limited protein synthesis capabilities. This adaptation underscores *T. pallidum*’s specialized metabolic strategies to thrive within host-associated, nutrient-limited environments. By providing a mechanistic view of these adaptations, the enzyme-constrained model ec-iTP251 identifies GPDH as a central component in maintaining redox balance, illustrating the critical role of alternative electron sinks in supporting energy conservation and survival. These findings highlight potential metabolic vulnerabilities in *T. pallidum*, suggesting that targeting GPDH and associated pathways could disrupt its redox stability and energy metabolism. Future work aimed at experimentally validating the role of GPDH could further assess its potential as a therapeutic target, offering new insights into strategies for weakening the bacterium’s resilience and pathogenicity within the host environment.

## MATERIALS AND METHODS

### Genome-Scale Metabolic Model Reconstruction and Refinement

We generated an initial draft of the genome-scale metabolic model for *T. pallidum* (RefSeq: NC_021490) using the KBase platform, where 3 reactions were added for gap filling purposes. Given the obligate host dependency and energy limitations of *T. pallidum*, extensive manual refinement was required to ensure that core metabolic pathways accurately reflected its unique physiology. Initial curation focused on central carbon metabolism, with adjustments to glycolytic and energy-conserving reactions. Specifically, the ATP-dependent phosphofructokinase (ATP-PFK) reaction was replaced with a pyrophosphate-dependent phosphofructokinase (PPi-PFK), reflecting *T. pallidum’s* reliance on PPi as an alternative to ATP (1). This metabolic adaptation, shared with related spirochetes like *Borrelia burgdorferi*, aligns with the bacterium’s strategy to conserve ATP in its energy-limited environment (48). Additionally, thirteen PTS reactions were removed, as experimental evidence confirms their absence in *T. pallidum’s* metabolic repertoire (49).

Targeted gap-filling, informed by high-resolution proteomic data, addressed additional gaps in the metabolic network. Eight reactions critical to *T. pallidum*’s central and peripheral metabolic pathways were added, encompassing nucleotide biosynthesis, lipid biosynthesis, amino acid metabolism, and cofactor and energy metabolism (see Table S4 for details). For nucleotide biosynthesis, reactions supporting pyrimidine and purine metabolism were incorporated to fulfill the demands of DNA replication and repair. Lipid biosynthesis pathways were expanded to include terpenoid backbone synthesis, reflecting *T. pallidum*’s requirements for membrane components essential for cellular integrity and function. Enhancements to amino acid metabolism ensured the synthesis of alanine, aspartate, and glutamate, supporting basic protein synthesis and cellular maintenance. Cofactor and energy metabolism pathways, including thiamine, nicotinate, and NAD⁺/NADH, were refined to maintain redox balance and energy production. One-carbon metabolism pathways were added to complete folate-related biosynthesis. Furthermore, D-lactate dehydrogenase was added to represent acetogenic ATP generation, enabling ATP synthesis independent of glycolysis.

Following these refinements, iTP251 was tested for its capacity to support growth on *T. pallidum*’s known carbon sources, including glucose, pyruvate, and mannose, to confirm biological accuracy. For pyruvate utilization, a transport reaction was added, while for mannose, both a transport reaction and a metabolic conversion from mannose-6-phosphate to fructose-6-phosphate were incorporated to enable substrate-specific growth.

### Biomass Composition and Maintenance Parameters

The biomass composition of iTP251 was adapted from the *Borrelia burgdorferi* (iBB151) metabolic model to approximate the cellular macromolecular makeup of *T. pallidum*(*30*). The biomass was defined as 70% protein, 20% lipid, and 5% carbohydrate on a dry weight basis, consistent with published physiological data of this bacteria (Table S1). To ensure accurate growth yield predictions, the biomass equation was standardized to achieve a molecular weight of 1 g/mmol.

Maintenance energy parameters, GAM and NGAM, were estimated to reflect *T. pallidum*’s unique physiology. Following Pirt’s equation was applied to calculate these parameters:

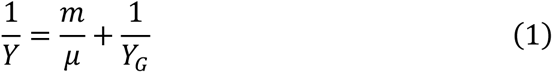

where *Y* is the observed growth yield, *Y_G_* is the maximum growth yield, *μ* is the specific growth rate, and *m* represents the maintenance coefficient. Theoretical parameters were initially derived from *Pseudomonas putida* due to its well-characterized physiology and were then adjusted to fit the metabolic characteristics of *T. pallidum* (50). Using the repurposed Prit’s equation for *T. pallidum*, we estimated GAM to be 48.69 mmol/gCDW/hr (Text S1, Table S2). This value was calibrated iteratively through flux balance simulations, ensuring agreement with its predicted growth characteristics. NGAM, representing baseline ATP requirements for essential cellular functions, was determined by simulating a zero-growth condition in the ec-iTP251 model. The glucose uptake rate (reaction “EX_cpd00027_e0”) was fixed at −0.0447 mmol/gDW/hr, representing minimal glucose consumption for non-growth maintenance. The model’s objective was set to maximize flux through the ATPase reaction (“rxn05145_c0”), which yielded an NGAM value of 1.50 mmol ATP/gDW.

Both GAM and NGAM values were subsequently incorporated as fixed parameters, ensuring accurate representation of maintenance energy across simulated environments.

### *In Silico* Reaction and Gene Essentiality Analysis

Reaction and gene essentiality analyses were conducted using the COBRApy toolbox to evaluate the predictive capacity of the iTP251 model. Reaction essentiality was determined by simulating single-reaction knockouts, where a reaction was classified as essential if its deletion reduced biomass flux by more than 90% compared to the wild-type model. Gene essentiality was assessed by simulating single-gene knockouts under the same criteria. Experimentally validated essentiality datasets for *Escherichia coli* and *Neisseria gonorrhoeae* were obtained from the Online Gene Essentiality (OGEE) database (51) for cross-species validation. Orthologous genes were identified using BLAST (52, 53), with an E-value threshold of 1e-5. This dual analysis provided a comprehensive evaluation of essential metabolic functions in *T. pallidum* while benchmarking the model’s predictions against established datasets for related organisms (Table S5).

### Enzyme-Constrained Model Development and Integration of Enzyme Kinetics

To enhance the physiological relevance of the iTP251 model, we developed ec-iTP251 by integrating *K_cat_* and molecular weights MW for reactions with GPR associations. The DLKcat tool predicted *K_cat_* values based on homology and structural properties for 391 out of the 471 GPR-associated reactions, thus providing estimates for reactions lacking direct experimental measurements (Table S6). Additionally, molecular weights of enzymes were calculated from amino acid sequences obtained from the KEGG database (54–56).

For the 80 reactions lacking *K_cat_* data, a Monte Carlo approach was employed to simulate biological variability. This method generated 100 sets of *K_cat_* values by stochastically sampling from distributions derived from the DLKcat predictions (Table S6). Each set was incorporated into the ec-iTP251 model, creating an ensemble of 100 models with diverse metabolic configurations.

To identify the most biologically relevant model, we calculated the total cellular protein demand (ϕ*_MC_*) for each configuration, defined as:

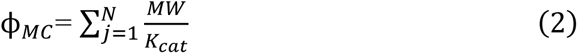

The ensemble model with the lowest total protein demand was selected as the most protein-efficient representation of *T. pallidum*’s metabolism.

### Validation Against Proteomic Data

To assess the accuracy of ec-iTP251 in predicting protein allocation, we validated the model against a high-resolution proteomic dataset. We first calculated the total protein associated with the central carbon pathway and then determined the fractional protein allocation required for each enzyme-associated reaction. Afterwards, we log-transformed these fractional values. This process was conducted for both the ec-iTP251 predictions and the experimental proteomics data, enabling a direct comparison between the two. Pearson’s correlation coefficient (*R*) was then applied to quantify the degree of agreement, and the coefficient of determination (*R*^2^) was calculated to measure predictive accuracy. The validation yielded an *R*^2^ value of 0.876, indicating a strong correlation between predicted and observed enzyme allocations. These results confirm the reliability of ec-iTP251 in accurately simulating protein distribution.

### Max-Min Driving Force Analysis

To evaluate the thermodynamic feasibility and compare the driving forces for lactate uptake versus lactate secretion, we applied MDF analysis. MDF identifies the minimal Gibbs free energy dissipation required to sustain flux directionality in a metabolic pathway, optimizing for the smallest driving force across all reactions. This approach is particularly useful in metabolic systems with limited metabolomic data, as it can be applied using plausible concentration ranges to ensure thermodynamically favorable reaction conditions.

The MDF formulation is outlined below:

*Maximize B*

*Subject to*

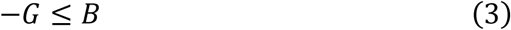

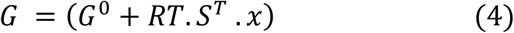

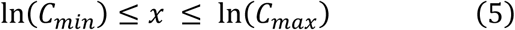

Here, *G*^0^ represents the standard Gibbs free energy obtained using the eQuilibrator API (57) *R* is the gas constant; *T* is the temperature (37°C); *x* is the vector of metabolite log-concentrations; and *C_min_* (1nM) and *C_max_* (10mM) define the lower and upper metabolite concentration bounds. The objective *B* represents the tightest lower bound on the driving force across all reactions, ensuring they operate as far from equilibrium as feasible within physiological constraints. Using this MDF framework, we conducted separate optimizations for both lactate uptake and lactate secretion conditions to capture and compare the thermodynamic driving force profiles.

### Substrate-Specific Metabolic Optimization under Protein Constraints

To analyze *T. pallidum*’s metabolic adaptations across the known carbon sources, substrate-specific optimizations were performed using the ec-iTP251 model. The objective was to identify reactions involved in ATP generation and NAD⁺ regeneration under protein-limited conditions.

For each substrate, specific constraints were implemented to account for its unique metabolic demands. For example, biomass production flux (*v_biomass_*) was fixed at 0.0231 *h*^−1^ and total cellular protein content (ϕ_FBA_) at 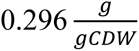 for glucose conditions. These values were readjusted for pyruvate and mannose to reflect their distinct metabolic profiles.

The optimization involved two steps. First, total protein allocation was minimized while satisfying biomass production requirements:

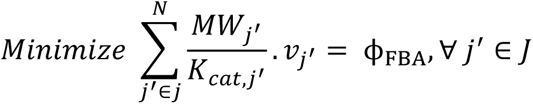

*Subject to*

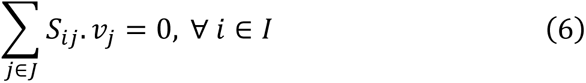

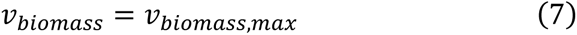

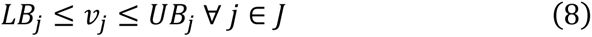

Here, *I* and *J* are the sets of metabolites and reactions in the model, respectively. *S_ij_* is the stoichiometric coefficient of metabolite *i* in reaction *j*, *j*^′^ is the set of reactions for which GPR is available, and *v_j_* is the flux value of reaction *j*. Parameters *LB_j_* and *UB_j_* denote the minimum and maximum allowable fluxes for reactions *j*, respectively.

In the second stage, parsimonious flux balance analysis (58) was performed with fixed ϕ_FBA_ and *v_biomass_*, minimizing enzyme usage while adhering to protein constraints. The resulting flux distributions were analyzed to identify substrate-specific differences in ATP generation and NAD⁺ regeneration. Key reactions contributing to these processes in Table S3.

### Software and Computational Tools

All computational analyses were performed using Python, specifically with the COBRApy 0.29.0 package for constraint-based modeling. The DLKcat tool was employed for predicting *K_cat_* values, and MEMOTE was used for assessing model quality and consistency. All simulations and analyses were conducted on a Windows 11 Enterprise operating system (Version 22H2), running on a machine equipped with a 12th Gen Intel® Core™ i7-12700H CPU at 2.30 GHz, 32.0 GB of RAM, and a 64-bit x64-based processor.

## DATA AVAILABILITY

The codes and resources supporting this study are available on GitHub at (https://github.com/ssbio/Treponema_pallidum_Nichols).

## ACKNOWLEDGMENTS

We gratefully acknowledge the funding support from the National Institute of Health (NIH) R35 MIRA grant (5R35GM143009) and National Science Foundation (NSF) CAREER grant (1943310).

## Notes

### Competing Interest Statement

The authors have declared no competing interest.

### Summary of Updates

Mathematical expressions were reformatted to address display issues.

